# SAMHD1 Impairs HIV-1 Gene Expression and Reactivation of Viral Latency in CD4^+^ T-cells

**DOI:** 10.1101/270843

**Authors:** Jenna M. Antonucci, Sun Hee Kim, Corine St. Gelais, Serena Bonifati, Olga Buzovetsky, Kirsten M. Knecht, Alice A. Duchon, Yong Xiong, Karin Musier-Forsyth, Li Wu

**Author notes:** Present address: HIV Dynamics and Replication Program, National Cancer Institute, Frederick, Maryland, USA. Address correspondence to: Li Wu,.

## Abstract

Sterile alpha motif and HD domain-containing protein 1 (SAMHD1) restricts human immunodeficiency virus type 1 (HIV-1) replication in non-dividing cells by degrading intracellular deoxynucleoside triphosphates (dNTPs). SAMHD1 is highly expressed in resting CD4^+^ T-cells that are important for the HIV-1 reservoir and viral latency; however, whether SAMHD1 affects HIV-1 latency is unknown. Recombinant SAMHD1 binds HIV-1 DNA or RNA fragments *in vitro*, but the function of this binding remains unclear. Here we investigate the effect of SAMHD1 on HIV-1 gene expression and reactivation of viral latency. We found that endogenous SAMHD1 impaired HIV-1 LTR activity in monocytic THP-1 cells and HIV-1 reactivation in latently infected primary CD4^+^ T-cells. Overexpression of wild-type (WT) SAMHD1 suppressed HIV-1 long terminal repeat (LTR)-driven gene expression at the level of transcription. SAMHD1 overexpression also suppressed LTR activity from human T-cell leukemia virus type 1 (HTLV-1), but not from murine leukemia virus (MLV), suggesting specific suppression of retroviral LTR-driven gene expression. WT SAMHD1 bound to proviral DNA and impaired reactivation of HIV-1 gene expression in latently infected J-Lat cells. In contrast, a nonphosphorylated mutant (T592A) and a dNTP triphosphohydrolase (dNTPase) inactive mutant (H206D/R207N, or HD/RN) of SAMHD1 failed to efficiently suppress HIV-1 LTR-driven gene expression and reactivation of latent virus. Purified recombinant WT SAMHD1, but not T592A and HD/RN mutants, bound to fragments of the HIV-1 LTR *in vitro*. These findings suggest that SAMHD1-mediated suppression of HIV-1 LTR-driven gene expression contributes to regulation of viral latency in CD4^+^ T-cells.

**IMPORTANCE:** A critical barrier to developing a cure for HIV-1 infection is the long-lived viral reservoir that exists in resting CD4^+^ T-cells, the main targets of HIV-1. The viral reservoir is maintained through a variety of mechanisms, including regulation of the HIV-1 LTR promoter. The host protein SAMHD1 restricts HIV-1 replication in non-dividing cells, but its role in HIV-1 latency remains unknown. Here we report a new function of SAMHD1 in regulating HIV-1 latency. We found that SAMHD1 suppressed HIV-1 LTR promoter-driven gene expression and reactivation of viral latency in cell lines and primary CD4^+^ T-cells. Furthermore, SAMHD1 bound to the HIV-1 LTR *in vitro* and in a latently infected CD4^+^ T-cell line, suggesting that the binding may negatively modulate reactivation of HIV-1 latency. Our findings indicate a novel role for SAMHD1 in regulating HIV-1 latency, which enhances our understanding of the mechanisms regulating proviral gene expression in CD4^+^ T-cells.

## INTRODUCTION

SAMHD1 is the only identified mammalian dNTPase (1, 2) with a well-characterized role in downregulation of intracellular dNTP levels (3), a mechanism by which SAMHD1 acts as a restriction factor against the infection of retroviruses (4, 5) and several DNA viruses (6–10) in non-dividing myeloid cells (11, 12) and quiescent CD4^+^ T-cells (13, 14). Additionally, *in vitro* studies indicate that SAMHD1 is a single-stranded (ss) nucleic acid (NA) binding protein (15–18), although the function of this binding activity in cells remains unknown. One report suggested that SAMHD1 uses its RNA binding potential to exert a ribonuclease activity against HIV-1 genomic RNA (19); however, recent studies do not support this observation (20–23).

SAMHD1 less efficiently restricts retroviral replication in dividing cells due to phosphorylation of SAMHD1 at Thr 592 (T592) (24–29). The dNTPase activity of SAMHD1 requires the catalytic H206 and D207 residues of the HD domain (30, 31). While mutations to either H206 or D207 abrogated ssDNA binding *in vitro* (15), the effect of nonphosphorylated T592 on ssDNA binding has not been described. The binding of ssNA occurs at the dimer-dimer interface on free monomers and dimers of SAMHD1. This interaction prevents the formation of catalytically active tetramers (18), suggesting a dynamic mechanism where SAMHD1 may regulate its potent dNTPase activity through NA binding. However, the effect of SAMHD1 and NA binding on HIV-1 infection or viral gene expression is unknown.

HIV-1 latency occurs post-integration when a proviral reservoir is formed within a population of resting memory CD4^+^ T-cells (32). By forming a stable reservoir and preventing immune clearance of infection, HIV-1 is able to persist in the host despite effective treatment with antiretroviral therapy (33). Although HIV-1 proviral DNA is transcriptionally silent in latently infected CD4^+^ T-cells, reactivation of intact provirus can result in the production of infectious virions (34, 35). There are several mechanisms that contribute to HIV-1 latency, including sequestration of host transcription factors in the cytoplasm and transcriptional repression (32, 35). The 5´ LTR promoter of HIV-1 proviral DNA contains several cellular transcription factor-binding sites, with transcription factors activated by external stimuli to enhance HIV-1 gene expression (36). Known cellular reservoirs of latent HIV-1 proviral DNA include quiescent CD4^+^ T-cells and macrophages (37–39). Although HIV-1 does not productively replicate in resting CD4^+^ T-cells, a stable state of latent infection does exist in these cells (40, 41). SAMHD1 blocks reverse transcription leading to HIV-1 restriction in resting CD4^+^ T-cells (13, 14); however, whether SAMHD1 affects reactivation of HIV-1 proviral DNA in latently infected CD4^+^ T-cells remains unknown.

In this study, we demonstrate that SAMHD1 suppresses HIV-1 LTR-driven gene expression and binds to the LTR promoter in a latently infected cell line model. Furthermore, endogenous SAMHD1 suppresses HIV-1 LTR-driven gene expression in a monocytic THP-1 cells and viral reactivation in latently infected primary CD4^+^ T-cells. Our findings suggest that SAMHD1-mediated suppression of HIV-1 gene expression contributes to regulation of viral latency in primary CD4^+^ T-cells, thereby identifying a novel role of SAMHD1 in modulating HIV-1 infection.

## RESULTS

### Exogenous SAMHD1 expression suppresses HIV-1 LTR-driven gene expression in HEK293T cells

Transcriptional activation of the HIV-1 provirus is regulated by interactions between the LTR promoter and several host and viral proteins (36). However, the effect of SAMHD1 expression on HIV-1 LTR-driven gene expression is unknown. To address this question, we performed an HIV-1 LTR-driven firefly luciferase (FF-luc) reporter assay using HEK293T cells. To examine transfection efficiency, a *Renilla* luciferase (Ren-luc) reporter driven by the herpes simplex virus (HSV) *thymidine kinase* (*TK*) promoter was used as a control (42). Expression of increasing levels of exogenous SAMHD1 did not change Ren-luc protein or mRNA expression (Fig. 1A-C), indicating comparable transfection efficiency among different samples, and that SAMHD1 overexpression did not affect *TK*-promoter driven gene expression. In contrast, when normalized with the Ren-luc control and compared to an empty vector, SAMHD1 expression resulted in 70-85% suppression of FF-luc activity (Fig. 1D) and *FF-luc* mRNA levels (Fig. 1E) in a dose-dependent manner. These data suggest that exogenous SAMHD1 expression suppresses HIV-1 LTR-driven gene expression at the level of gene transcription.

**FIG 1.**
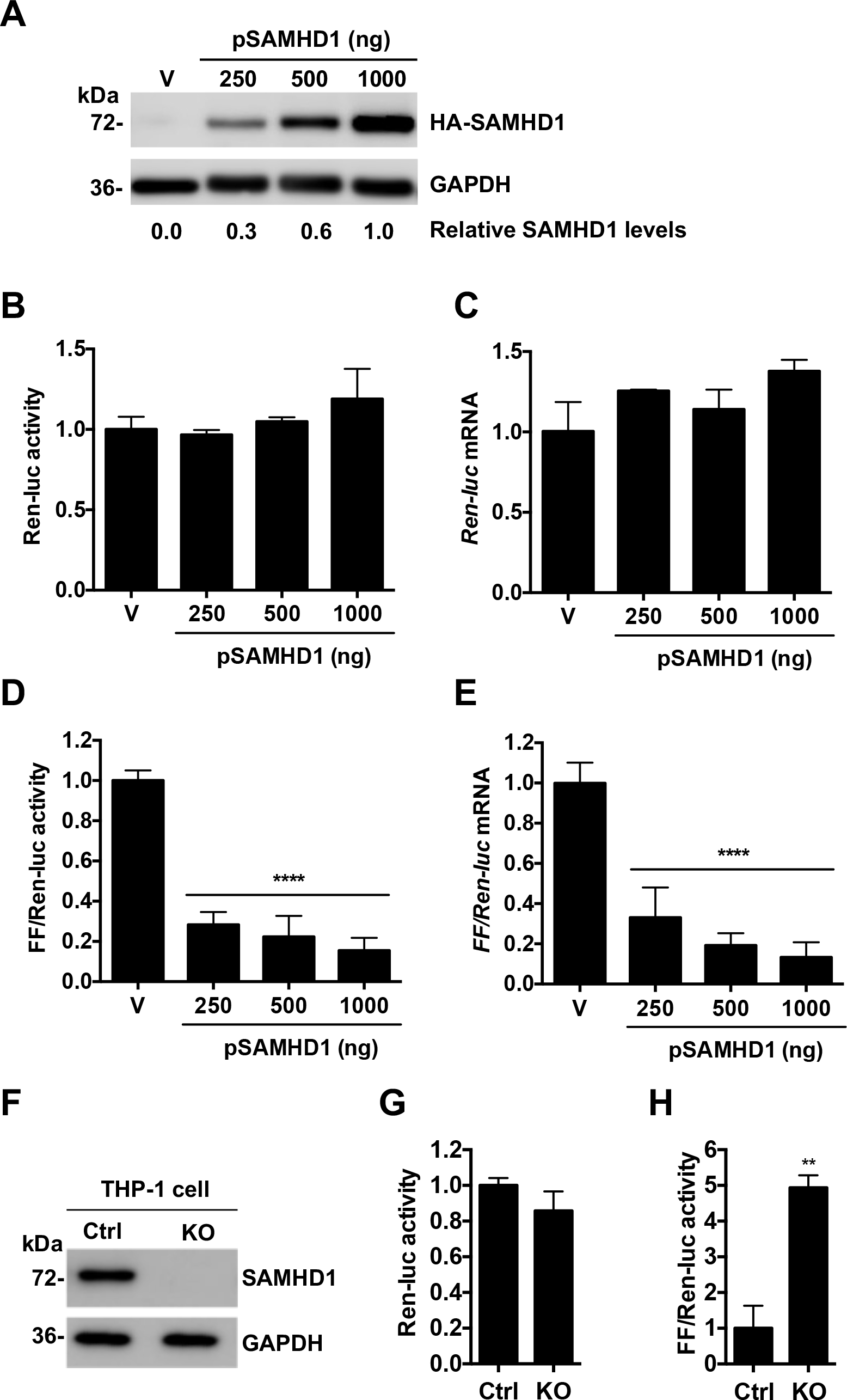
SAMHD1 suppresses HIV-1 LTR-driven luciferase expression. **(A-E)** An HIV-1 LTR-driven firefly luciferase (FF-Luc) construct was co-transfected with an empty vector (V) or increasing amounts of a plasmid encoding HA-tagged SAMHD1 (pSAMHD1) into HEK293T cells. Co-transfection of a construct encoding HSV TK-driven Renilla luciferase (Ren-Luc) was used as a control of transfection efficiency. **(A)** Overexpression of SAMHD1 was confirmed by immunoblotting. GAPDH was used as a loading control. Quantification of relative SAMHD1 expression levels by densitometry was normalized to GAPDH, with the level of 1000 ng pSAMHD1 sample set as 1. Ren-luc activity **(B)** and mRNA **(C)**, and FF-luc activity **(D)** and mRNA **(E)** were measured at 24 h post-transfection. **(B)** Ren-luc activity was normalized to total protein concentration. **(D and E)** FF-luc activity and mRNA levels were normalized to Ren-luc activity and mRNA levels, with vector levels set as 1. **(B-E)** Error bars show standard deviation (SD) of at least three independent experiments as analyzed by one-way ANOVA with Dunnett’s multiple comparison post test. ****, p≤0.0001 compared to vector control cells. **(F-H)** FF-luc and Ren-luc constructs were expressed by nucleofection in THP-1 control (Ctrl) cells or SAMHD1 knockout (KO) cells. SAMHD1 KO was confirmed by immunoblotting, with GAPDH used as a loading control **(F)**. Luciferase activity was measured at 48 h post-transduction, and raw Ren-luc values were normalized to total protein **(G)**, and FF-luc activity normalized to Ren-luc activity **(H)**. **, p≤0.01 compared to control cells.

### SAMHD1 silencing in THP-1 cells enhances HIV-1 LTR-driven gene expression

To determine whether endogenous SAMHD1 can suppress LTR-driven gene expression in cells, we performed the HIV-1 LTR reporter assay using human monocytic THP-1 cells expressing a high level of endogenous SAMHD1 (Ctrl) and SAMHD1 knockout (KO) (29). THP-1 control or KO cells were nucleofected with plasmids expressing FF-luc and Ren-luc. The lack of SAMHD1 expression in THP-1 KO cells was confirmed by immunoblotting (Fig. 1F). Consistent with the results from HEK293T cells, the Ren-luc activity was unchanged in THP-1 control and KO cells, confirming comparable transfection (Fig. 1G). When normalized with Ren-luc activity, KO cells showed a 4.5-fold increase of FF-Luc activity compared to control cells (Fig. 1H), indicating that endogenous SAMHD1 impairs HIV-1 LTR-driven gene expression in THP-1 cells.

### SAMHD1 suppresses gene expression driven by the LTR from HIV-1 and HTLV-1, but not from MLV

To examine the specificity of SAMHD1-mediated suppression of LTR-driven gene expression, we tested luciferase reporters driven by LTR promoters derived from HTLV-1 or MLV LTR in addition to the HIV-1 LTR FF-luc reporter. The *TK* promoter-driven Ren-luc reporter was used as a transfection control. As HIV-1 and HTLV-1 utilize viral proteins Tat (43) and Tax (44) respectively, to enhance viral transcription via transactivation, we compared the ability of SAMHD1 to suppress HIV-1 and HTLV-1 LTR-driven gene expression with or without transactivation. Increased levels of exogenous SAMHD1 expression in transfected HEK293T cells were confirmed by immunoblotting (Fig. 2A, C, E). Although SAMHD1 suppressed HIV-1 LTR-driven gene expression in the absence of Tat (Fig. 2B, black bars), Tat expression led to a 20 to 28-fold enhancement of HIV-1 LTR activity that was not suppressed by SAMHD1 (Fig. 2B, gray bars). Conversely, HTLV-1-LTR activity was potently suppressed by SAMHD1 expression, with up to 75% reduction in luciferase activity at the highest level of SAMHD1 expression in the presence or absence of Tax (Fig. 2D). SAMHD1 expression had no effect on MLV-LTR activity (Fig. 2F). These data suggest that SAMHD1 selectively suppresses retroviral LTR-driven gene expression.

**FIG 2.**
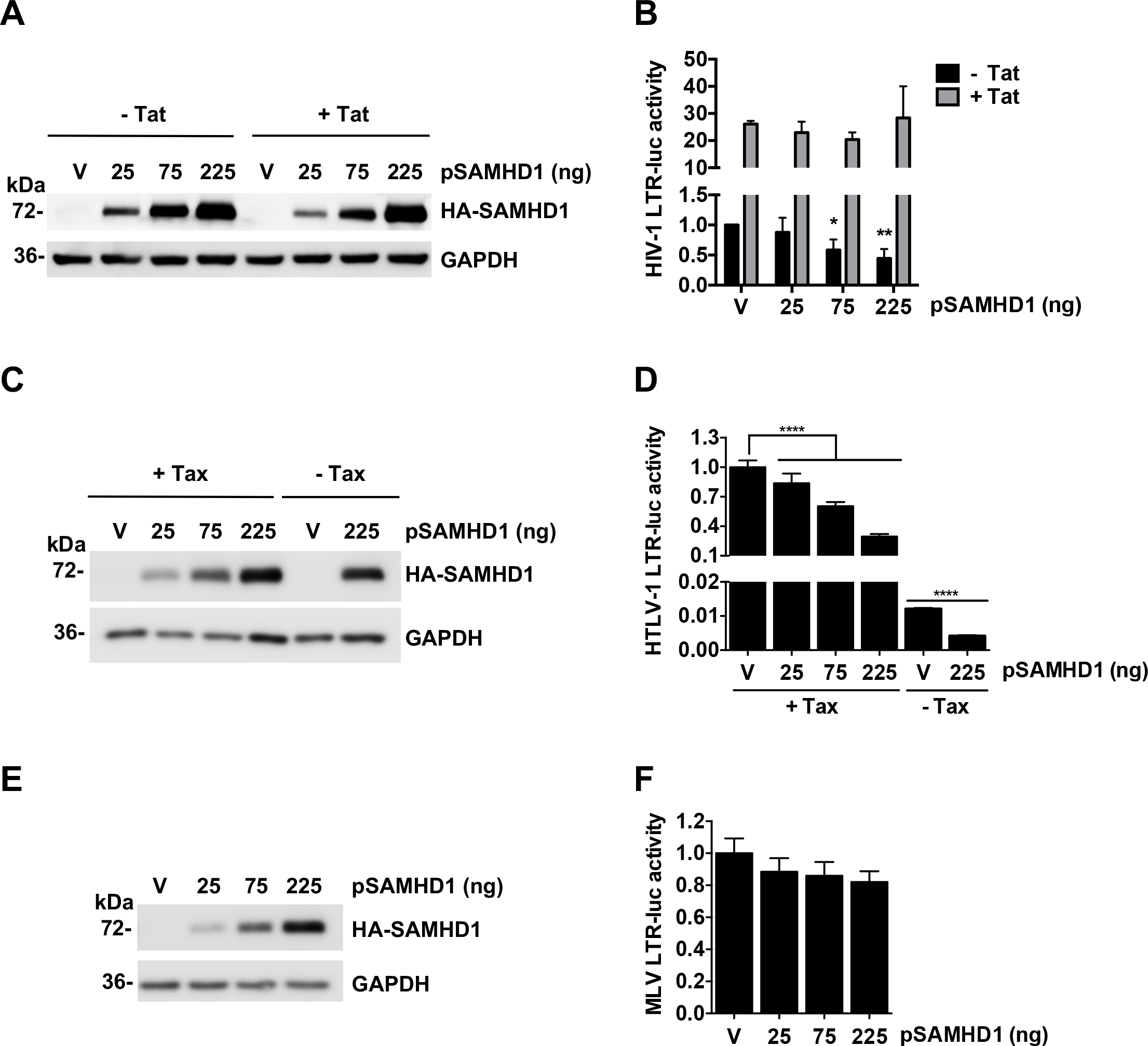
SAMHD1 suppresses gene expression driven by the LTR from HIV-1 and HTLV-1, but not from MLV. **(A-F)** HEK293T cells were transfected with an empty vector (V) or increasing amounts of constructs expressing HA-tagged SAMHD1 and either an HIV-1 LTR-driven FF-luc construct with or without HIV-1 Tat-expressing plasmid **(A-B)**, HTLV-1 LTR-driven FF-luc construct with or without HTLV-1 Tax-encoding plasmid **(C-D)**, or MLV LTR-driven FF-luc construct **(E-F)**. Overexpression of SAMHD1 was analyzed by immunoblotting **(A, C, E)** with GAPDH as a loading control. Co-transfection of Ren-luc was used as a control of transfection efficiency, with LTR-driven FF-luc activity normalized to Ren-luc activity. **(B, D, F)**. Luciferase activity was determined 24 h post transfection. Error bars show standard error mean (SEM) of three (HIV-luc +/− tat, HTLV-luc +tat) or two (HTLV-luc -tat, MLV-luc) independent experiments. Statistical analysis was performed by one-way ANOVA with Dunnett’s multiple comparison post test. *, p≤0.05, **, p≤0.01, and ****, p≤0.0001, compared to vector (V) control cells.

### Nonphosphorylated and dNTPase-inactive SAMHD1 mutants have impaired suppression of HIV-1 LTR activity

SAMHD1 is predominantly phosphorylated in HEK293T cells (25, 27). To assess the effect of dNTPase activity and T592 phosphorylation of SAMHD1 on suppression of LTR-driven gene expression, we tested a catalytically inactive SAMHD1 mutant HD/RN (30) and a nonphosphorylated T592A mutant (26). HEK293T cells were transfected with increasing amounts of plasmids encoding WT, T592A, or HD/RN mutant SAMHD1, along with the HIV-1 LTR-driven FF-luc reporter. Comparable WT and mutant SAMHD1 expression was confirmed by immunoblotting (Fig. 3A). Undetectable phosphorylation of the T592A mutant was confirmed in our previous studies (21). The *TK* promoter-driven Ren-luc reporter showed similar activity across all samples (Fig. 3B), confirming comparable transfection efficiency. Compared to the vector control and normalized with Ren-luc activity, WT SAMHD1 suppressed HIV-1 LTR-driven FF-luc expression up to 60% in a dose-dependent manner (Fig. 3C). In contrast to WT SAMHD1, low amounts of HD/RN or T592A mutants (125 and 250 ng plasmid input) did not significantly inhibit HIV-1 LTR activity. However, a modest inhibition of LTR-driven FF-luc activity (between 25-31%) was observed at the highest levels of mutant SAMHD1 expression (500 ng plasmid input) (Fig. 3C). These results indicated that T592A and HD/RN mutants have a diminished ability to suppress HIV-1 LTR-driven gene expression compared to WT SAMHD1, suggesting that SAMHD1-mediated suppression of HIV-1 gene expression is partially dependent on its dNTPase activity and T592 phosphorylation.

**FIG 3.**
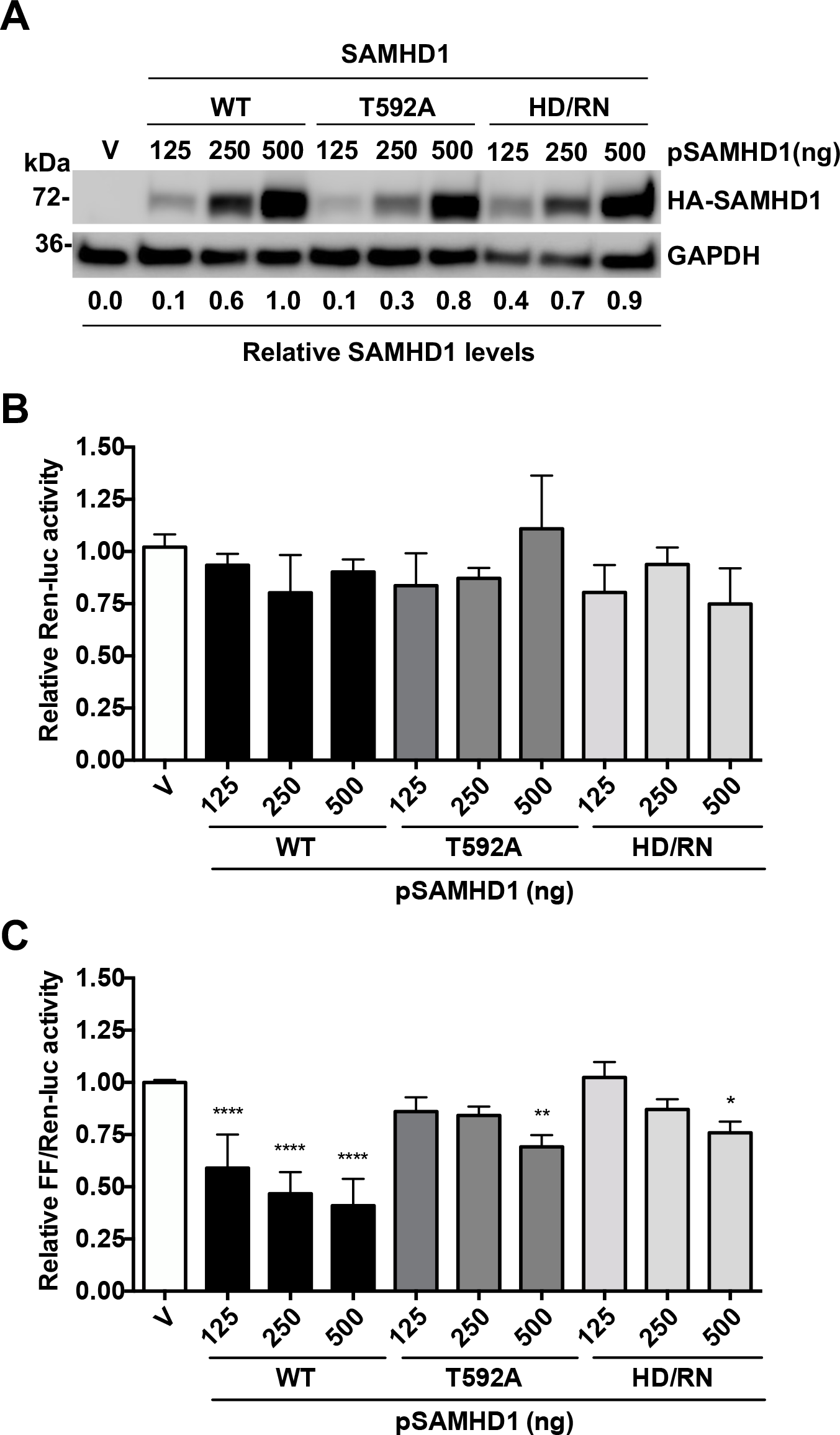
Nonphosphorylated and dNTPase-inactive SAMHD1 mutants have impaired suppression of HIV-1 LTR activity. **(A-C)** An HIV-1 LTR-driven FF-luc construct was cotransfected with increasing amounts of plasmids encoding HA-tagged WT, nonphosphorylated T592A, or dNTPase-inactive HD/RN mutant SAMHD1 into HEK293T cells. Co-transfection of Ren-luc was used as a control of transfection efficiency. **(A)** SAMHD1 expression was confirmed by immunoblotting. GAPDH was used as a loading control. Quantification of relative SAMHD1 expression levels by densitometry was normalized to GAPDH. Relative Ren-luc units were normalized to total protein concentration **(B)**, and relative FF-luc units were normalized to Ren-luc levels **(C)**. Vector cell luciferase activity set as 1. Statistical analysis was performed by one-way ANOVA with Dunnett’s multiple comparison post test. Error bars show SD of at least three independent experiments. *, p≤0.05, **, p≤0.01, and ****, p≤0.0001 compared to vector control cells.

### WT SAMHD1 impairs HIV-1 reactivation in latently infected J-Lat cells

To investigate whether SAMHD1 suppresses HIV-1 gene expression in CD4^+^ T-cells, we used Jurkat CD4^+^ T cell line-derived J-Lat cells (45). J-Lat cells have been used as an HIV-1 latency model, as they contain a full-length HIV-1 provirus with a *green fluorescent protein* (*gfp*) gene inserted in the *nef* region (45). Treatment of J-Lat cells with latency reversing agents (LRAs) results in activation of LTR-driven gene expression, indicated by an increase in GFP expression (45, 46). As J-Lat cells do not express detectable endogenous SAMHD1 protein, likely due to gene promoter methylation as reported in Jurkat cells (47), we stably expressed WT SAMHD1 in J-Lat cells by lentiviral transduction. Empty vector-transduced cells were used as a control. Previous studies showed that efficient SAMHD1 expression driven by the cytomegalovirus (CMV) immediate-early promoter of stably integrated lentiviral vector in monocytic cell lines is dependent on treatment of cells with phorbol 12-myristate 13-acetate (PMA) (4, 27), which is a protein C kinase agonist that activates the NF-κB signaling pathway (48, 49).

To activate HIV-1 gene expression in J-Lat cells we applied two LRAs, tumor necrosis factor alpha (TNFα) which induces HIV-1 gene expression by activating the NF-κB pathway (46, 50, 51), and PMA in conjunction with ionomycin (PMA+i) that has been shown to be the strongest activator of HIV-1 gene expression in several J-Lat cell clones (46). Treatment of J-Lat cells with TNFα (Fig. 4A) resulted in GFP expression in 38% of vector control cells (Fig. 4B), consistent with published data (45, 46). In contrast, expression of WT SAMHD1 reduced TNFα-induced GFP expression to 27% (Fig. 4B), suggesting that SAMHD1 impairs TNFα-induced HIV-1 reactivation. Treatment of WT SAMHD1-expressing J-Lat cells with PMA+i resulted in a significant increase in SAMHD1 expression (Fig. 4A) and a significant decrease in GFP-expression by 20% compared to vector control cells (Fig. 4B). While PMA+i treatment resulted in a 1.5-fold reduction of GFP mean fluorescence intensity (MFI) of SAMHD1-expressing cells compared to vector, the MFI of TNFα-treated cells was not significantly reduced by SAMHD1 expression (Fig. 4B).

**FIG 4.**
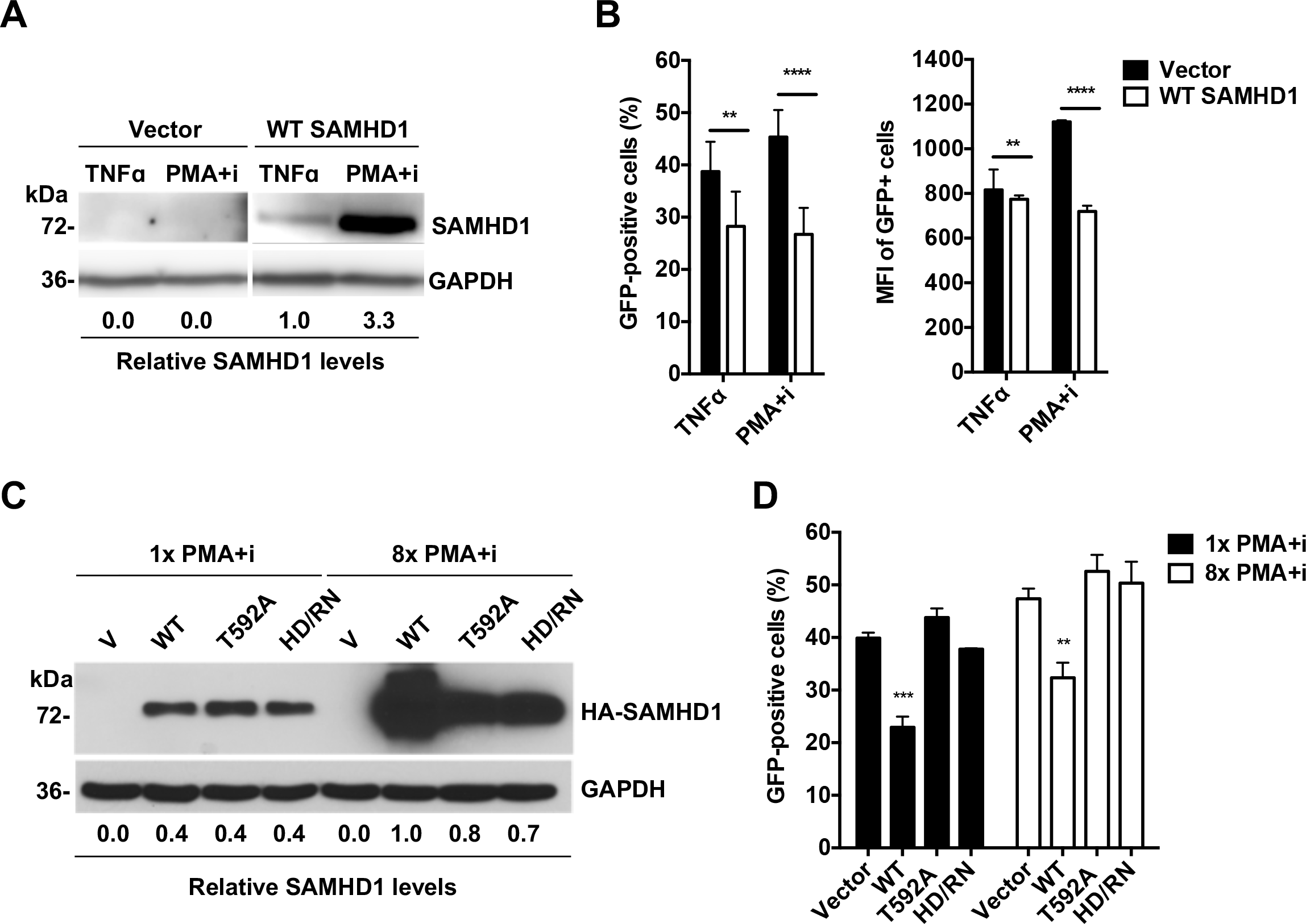
WT SAMHD1 impairs HIV-1 reactivation in latently infected J-Lat cells. **(A, C)** HA-tagged SAMHD1 WT or mutants, or an empty vector were stably expressed in J-Lat cells by lentiviral transduction. Quantification of relative SAMHD1 expression levels by densitometry was normalized to GAPDH. **(A-B)** The cells were treated with either 10 ng/mL TNFα, or 32 nM PMA with 1 μM ionomycin (PMA+i). At 24 h post-treatment, the expression of SAMHD1 was detected by immunoblotting, with quantification of SAMHD1 expression by densitometry normalized to GAPDH levels **(A)**. The percentage of GFP-positive cells and the relative GFP mean fluorescence intensity (MFI) were determined by flow cytometry **(B).** J-Lat cells expressing T592A and HD/RN mutants were treated with 1× or 8× PMA+i (1× corresponds to 16 nM PMA and 0.5 μM ionomycin), with expression of SAMHD1 measured and quantified by immunoblotting **(C)**. Latency reversal, as measured by percentage of GFP-positive cell population and MFI of GFP-positive cells, was determined by flow cytometry **(D).** Error bars in **(B and D)** represent SD from at least three independent experiments analyzed by two-way ANOVA and Dunnett’s multiple comparisons test. **, p≤0.01, ***, p≤0.001, and ****, p≤0.0001 (compared to vector cells in panels B and D).

To examine whether increased SAMHD1 expression in J-Lat cells could more efficiently suppress HIV-1 reactivation, we compared J-Lat cells treated with two PMA+i concentrations with a 8-fold difference (Fig. 4C and 4D). To examine whether the dNTPase activity or T592 phosphorylation of SAMHD1 affects its suppression of HIV-1 reactivation in J-Lat cells, we performed the analysis in J-Lat cells stably expressing WT SAMHD1, T592A, or HD/RN mutant by lentiviral transduction. Similar expression levels of WT SAMHD1 and the mutants were observed in 1× PMA+i-treated cells, while 8× PMA+i treatment highly increased the expression levels of WT SAMHD1, and mutant SAMHD1 to a lesser degree (Fig. 4C). WT SAMHD1-expressing cells had a 15% lower GFP-positive cell population compared to vector control cells at 1× PMA+i; however, this was not further enhanced with increased WT SAMHD1 expression at 8× PMA+i (Fig. 4D). While WT SAMHD1 suppressed HIV-1 reactivation at both 1× and 8× PMA+i treatment, neither T592A nor HD/RN mutant had a suppression effect (Fig. 4D), suggesting that T592 phosphorylation and dNTPase activity of SAMHD1 are likely involved in reactivation of HIV-1 latency.

### WT SAMHD1 binds to HIV-1 LTR of proviral DNA in J-Lat cells

One common mechanism by which host proteins modulate HIV-1 LTR activity is transcriptional repression by directly binding to the promoter (52). SAMHD1 is a DNA binding protein (20) capable of interacting with *in vitro* transcribed HIV-1 *gag* DNA fragments (15). However, the interaction between SAMHD1 and integrated HIV-1 proviral DNA in cells has not been reported. To address this question, we performed a chromatin immunoprecipitation coupled with quantitative real-time PCR (ChIP-qPCR) experiment in J-Lat cells expressing WT, T592A, or HD/RN SAMHD1. To induce high levels of SAMHD1 for efficient immunoprecipitation (IP), we treated the cells with increased PMA+i concentrations. Treatment with 8× PMA+i allowed for maximum SAMHD1 expression without cell death (data not shown). However, WT SAMHD1 expressed 20-30% greater than mutants under this condition (Fig. 4C). We treated the WT SAMHD1-expressing cells with 50% less PMA+i compared to that used for mutant-expressing cells, and obtained comparable levels of SAMHD1 (Fig. 5A). HIV-1 reactivation in all cell lines was measured by GFP expression (Fig. 5B). The WT SAMHD1-expressing J-Lat cells had a 17% lower GFP-positive population compared to vector control, T592A, and HD/RN-expressing cells, which was reflected in a 1.6-fold lower MFI (Fig. 5B). After IP of WT or mutant SAMHD1 from cells treated with PMA+i (Fig. 5A), total bound DNA was eluted and quantified by qPCR. We used PCR primers specific for different regions in the HIV-1 genome, including the LTR, *gag, vpr,* and *rev* genes, to characterize the regions of interaction between SAMHD1 and proviral DNA (Fig. 5C and Table 1). We also included *gfp*-specific PCR primers as an additional control, as *gfp* is a non-viral gene inserted in the *nef* gene of HIV-1 in J-Lat cells (45). We observed that only DNA fragments derived from the LTR (12% of input) bound to WT SAMHD1 (Fig. 5D). WT SAMHD1 did not bind other HIV-1 genes tested or the *gfp* gene. These data suggest that the SAMHD1-DNA interaction occurs in the LTR promoter region of HIV-1 provirus. Interestingly, analysis of the DNA eluted from IP products of T592A and HD/RN SAMHD1 revealed that neither mutant bound to tested HIV-1 DNA sequences or *gfp* cDNA (Fig. 5D). Taken together, these data indicate that mutant SAMHD1 cannot suppress latency reactivation or bind to proviral DNA, suggesting that direct binding to the HIV-1 LTR is partially responsible for the mechanism of SAMHD1-mediated suppression of LTR-driven gene expression.

**FIG 5.**
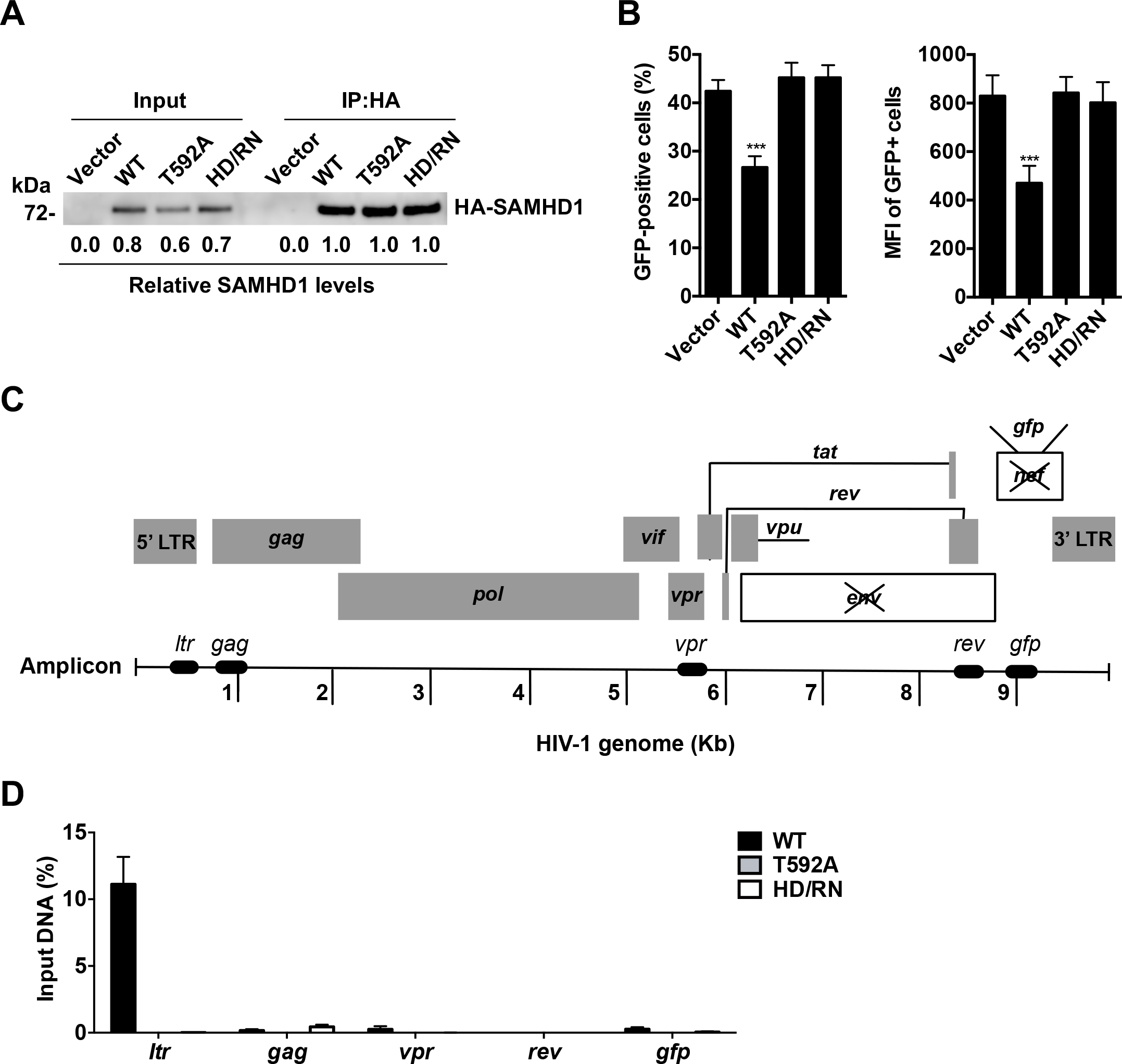
WT SAMHD1 binds to HIV-1 proviral DNA in latently infected J-Lat cells. (**A-D**) J-Lat cells were seeded in the presence of PMA+i for 24 h. To achieve similar expression levels of SAMHD1, WT SAMHD1-expressing cells were treated with 4× PMA+i, while vector control, T592A-, and HD/RN-expressing cells were treated with 8× PMA+i. SAMHD1 expression in input and IP lysates was analyzed by immunoblotting and densitometry analysis **(A)**. Latency reversal, as measured by percentage of GFP-positive cell population and MFI of GFP-positive cells, was determined by flow cytometry **(B)**. Three independent experiments were analyzed by two-way ANOVA and Dunnett’s multiple comparisons test, with error bars in **(B)** representing SEM. ***, p≤0.001 compared to vector cells. **(C)** Diagram of the location of the qPCR amplicons. Quantitative PCR data were normalized to spliced GAPDH levels and presented as percent of input in SAMHD1-expressing cells over vector cells **(D)**. Error bars in **(D)** represent SEM from two independent experiments.

**Table 1.**
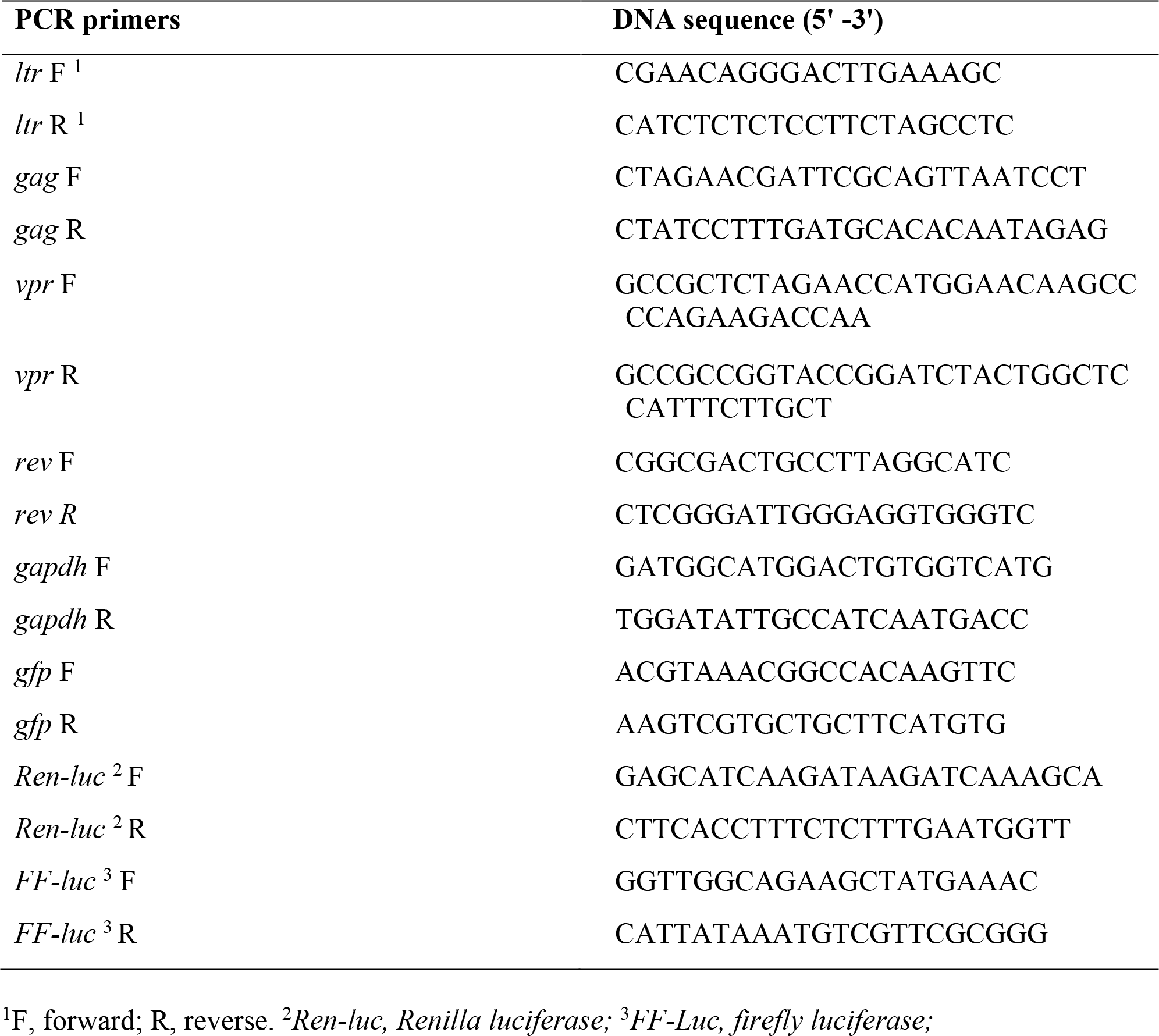
PCR primer sequences

### Purified recombinant WT SAMHD1 binds to HIV-1 LTR fragments *in vitro*

To investigate the underlying mechanism of SAMHD1-mediated suppression of HIV-1 LTR activity and viral reactivation in cells, we determined whether this suppression effect correlated with SAMHD1 binding to HIV-1 LTR *in vitro*. Fluorescence anisotropy (FA) (53) was used to measure the binding of WT SAMHD1, T592A or HD/RN mutant to a 90-mer 5’-6-carboxyfluorescein (6-FAM)-labeled DNA oligonucleotide derived from the HIV-1 LTR (Table 2). Binding was measured over a range of SAMHD1 concentrations and three monovalent ion concentrations (50, 100, and 150 mM) to determine whether the interaction is mediated by electrostatic interactions. While WT SAMHD1 binding to the HIV-1 LTR fragment was detected at all three salt concentrations tested, higher salt reduced the observed binding (Fig. 6A-C), which suggests that the interaction is mediated, at least in part, by electrostatic contacts. In contrast, no significant binding was observed for the T592A and HD/RN mutants even at the highest protein concentration (8,300 nM) (Fig. 6A-C). For WT SAMHD1, saturated or near-saturated binding was observed at 50 and 100 mM monovalent ions (Fig. 6A and 6B) with calculated apparent K_d_ values of 93 ± 8 and 242 ± 51 nM, respectively. Importantly, none of the SAMHD1 proteins bound to a 90-mer 6-FAM-labeled DNA oligonucleotide derived from a scrambled sequence of the HIV-1 LTR (Table 2), even at low (50 mM) monovalent ion concentration (Fig. 6D). These data indicate that WT SAMHD1 binds specifically to HIV-1 LTR-derived fragments in a salt sensitive manner.

**FIG 6.**
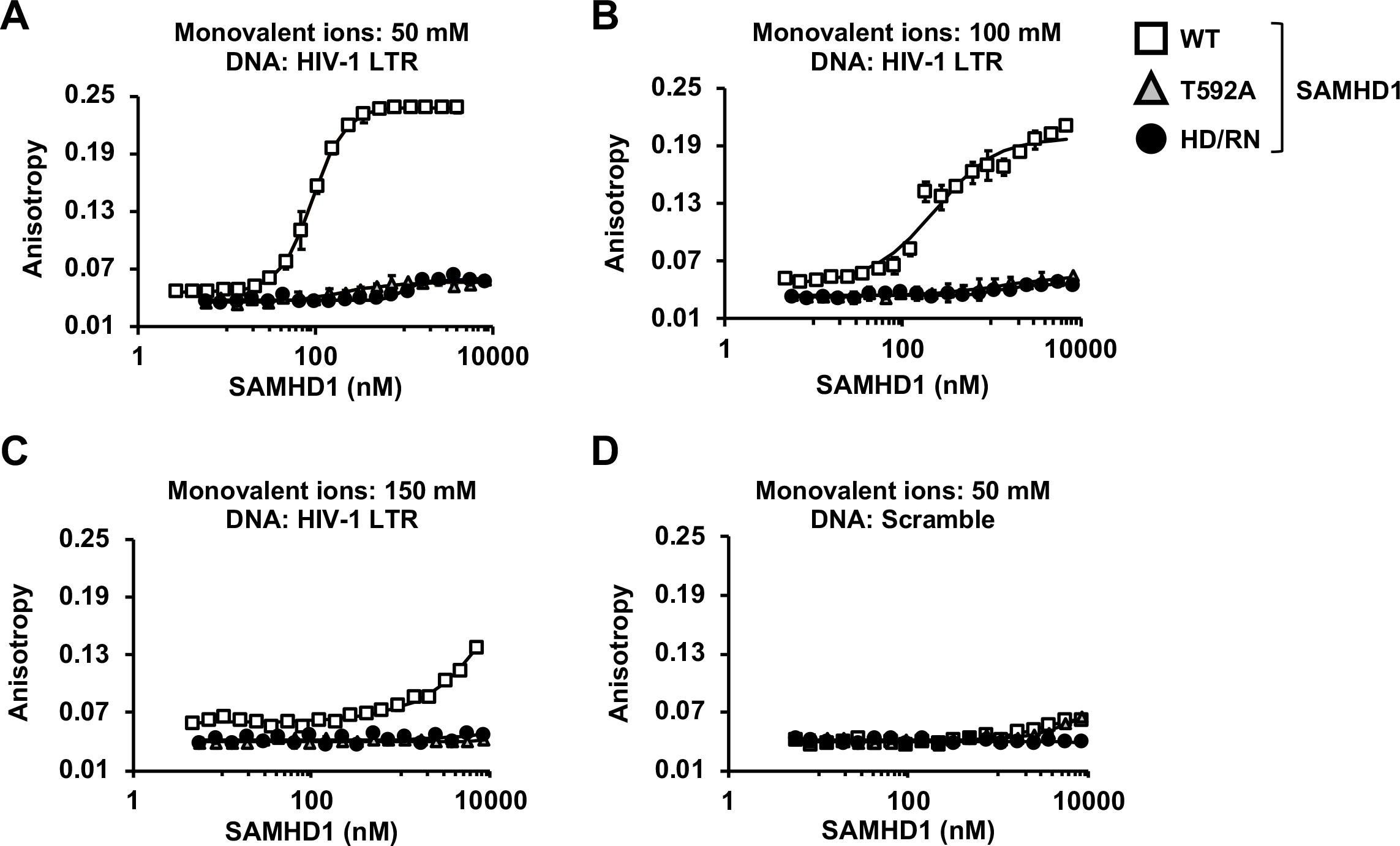
Specific binding of WT SAMHD1 to an HIV-1 LTR fragment *in vitro*. **(A-D)** Results of FA binding assays for WT, T592A or HD/RN mutant SAMHD1 binding to a 90-mer fragment of the HIV-1 LTR in 50 mM, 100 mM, or 150 mM monovalent ions (25, 50, or 75 mM of each NaCl and KCl) **(A-C,** respectively). Binding to a 90-mer scrambled DNA oligonucleotide was also tested at 50 mM monovalent ions (25 mM of each NaCl and KCl) **(D)**. Error bars indicate the SD from three independent experiments.

**Table 2.**
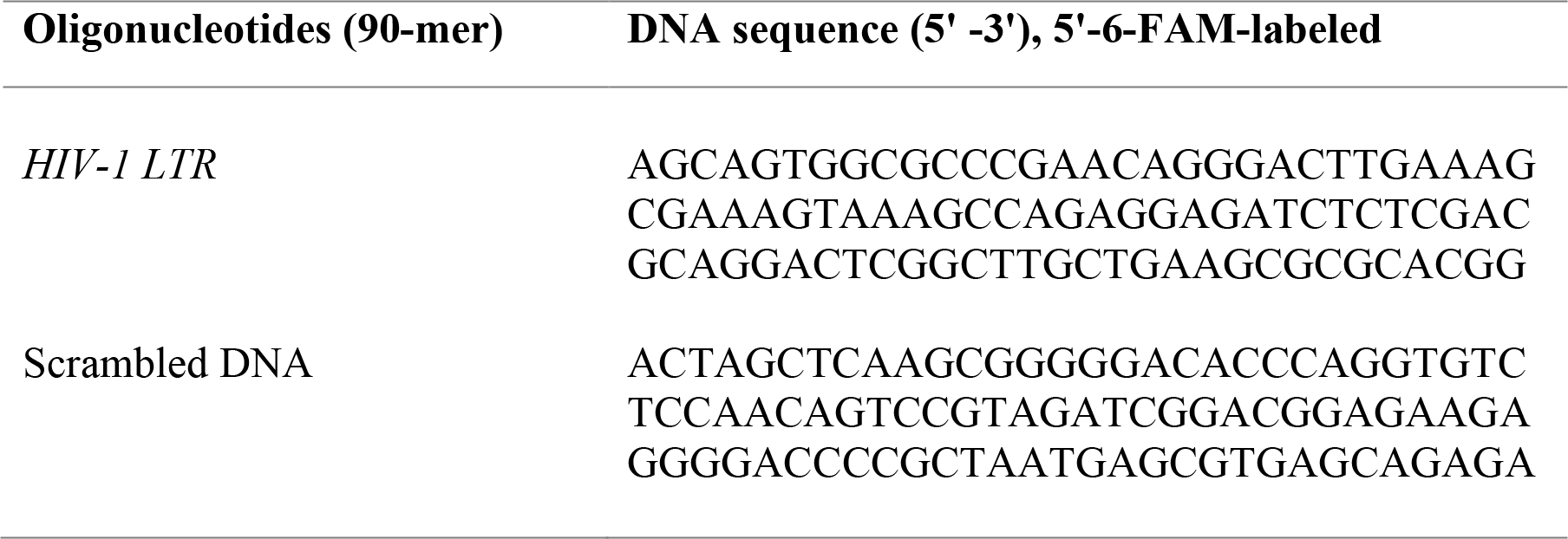
Sequences of oligonucleotides used in anisotropy binding assays

### SAMHD1 knockdown promotes HIV-1 reactivation in latently infected primary CD4^+^ T-cells

To examine the effect of endogenous SAMHD1 on HIV-1 reactivation in primary CD4^+^ T-cells, we utilized the established central memory T-cell (T_CM_) model of HIV-1 latency (40, 54) as a depicted in the protocol in Fig. 7A. Using a *SAMHD1*-specific shRNA and established method (54, 55), we knocked down 40-50% of endogenous SAMHD1 expression in latently infected T_CM_ derived from naïve CD4^+^ T-cells isolated from three healthy donors (Fig. 7B). Next, we activated latently infected GFP-reporter HIV-1 in T_CM_ by CD3/CD28 antibody treatment and measured GFP expression as a readout of latency reactivation. As a negative control, T_CM_ treated with media did not express GFP (1% background). Upon activation of latently infected T_CM_, partial knockdown of SAMHD1 enhanced HIV-1 reactivation by 1.6-fold compared to cells transduced with an empty shRNA vector, as measured by % GFP-positive cell population and MFI of GFP-positive cells (Fig. 7B and 7C). These results confirm that endogenous SAMHD1 acts as a negative regulator of HIV-1 reactivation in latently infected primary CD4^+^ T-cells.

**FIG 7.**
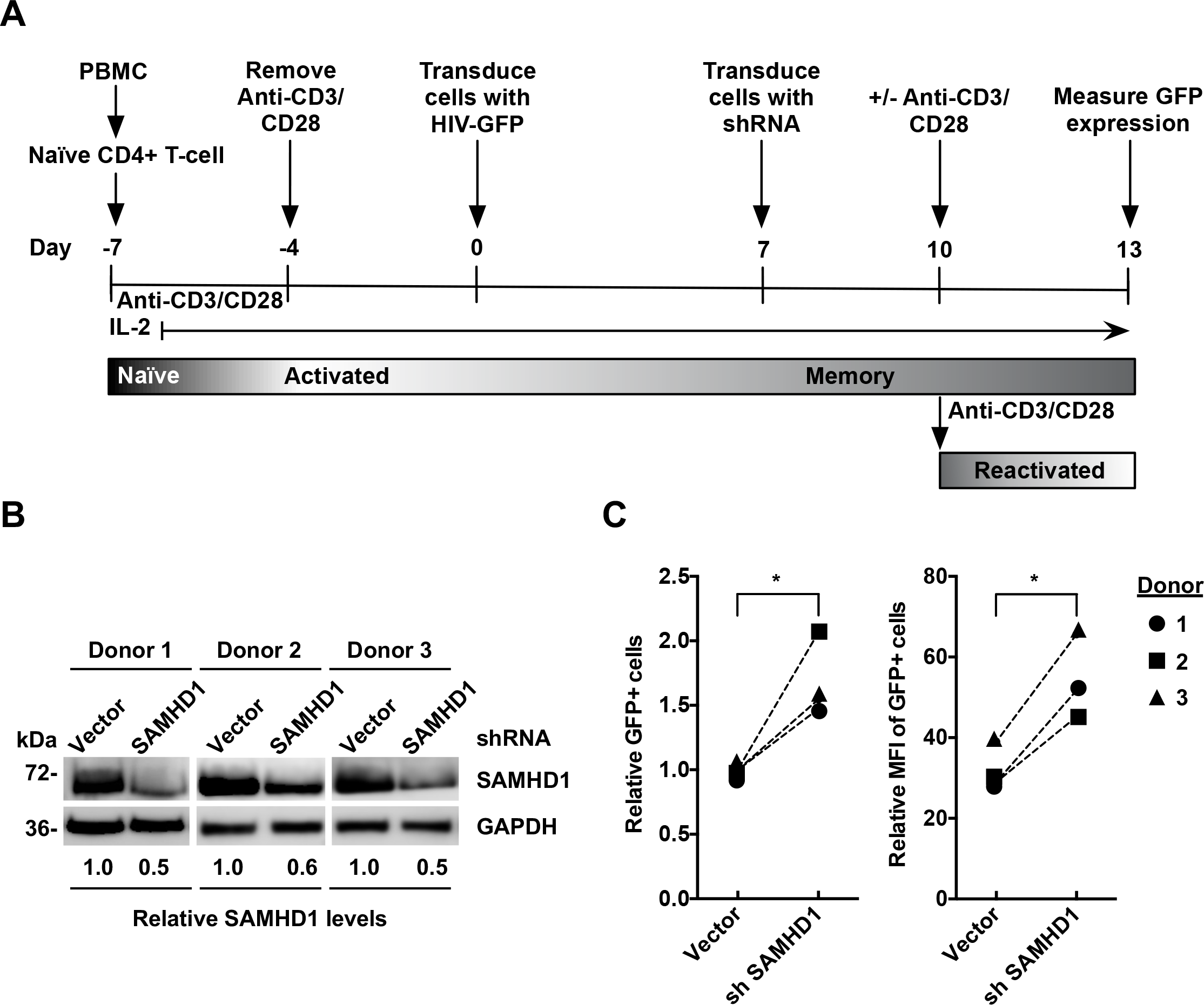
Endogenous SAMHD1 impairs HIV-1 reactivation in latently infected primary CD4^+^ T-cells. **(A)** Protocol summary. Naïve CD4^+^ T-cells were isolated from PBMCs from three different healthy donors, activated by incubation with anti-CD3/CD28 antibody-coated beads, and transduced with a single-cycle HIV-1 containing a *GFP* reporter (HIV-GFP) to produce a primary T_CM_ cell model of latency. After infection and culture to produce latently infected quiescent CD4^+^ T-cells, transduction with either empty vector or lentiviral vectors containing SAMHD1-specific shRNA to knockdown of SAMHD1. SAMHD1 expression was measured by immunoblotting and GAPDH was a loading control **(B)**. After stimulation of the cells with anti-CD3/CD28, HIV-1 reactivation was measured by flow cytometry. Relative changes in the GFP-positive cell population and MFI of GFP-positive cells were quantified **(C)**, with each line indicating the result from one donor. *, p≤ 0.05.

## DISCUSSION

One of the hallmarks of HIV-1 persistence is the maintenance of a long-lived stable proviral reservoir that is formed after infection in resting CD4^+^ T-cells (32, 56). Although the integrated provirus is transcriptionally silent, it is capable of full reactivation and production of infectious virus upon discontinuation of therapy or treatment with LRAs (32, 41, 56). In this study, we tested the hypothesis that SAMHD1 plays a role in negatively regulating HIV-1 reactivation and viral latency by suppressing HIV-1 LTR-driven gene expression.

We demonstrated that WT SAMHD1 suppresses HIV-1 LTR-driven gene expression, in the absence of Tat, in HEK293T and THP-1 cells. Previous work confirmed that SAMHD1 does not degrade HIV-1 genomic RNA or mRNA (20, 21), thereby excluding the possibility that mRNA degradation causes the suppression. We observed that SAMHD1 potently suppressed the HTLV-1 LTR independently of Tax expression but had no effect on the MLV LTR or HIV-1 LTR in the presence of Tat. These differences in suppression could be the result of variations in transcriptional control of each LTR. Tat transactivation of the HIV-1 LTR occurs through direct binding of Tat to the HIV-1 transactivation-responsive region (43, 57). Tat transactivation may saturate LTR activity and mask a SAMHD1-mediated suppressive effect. The Tat-TAR binding affinity is particularly tight, with a K_d_ of 1-3 nM, making effective competition by SAMHD1 very unlikely (43, 58). Moreover, we previously observed that SAMHD1 expression does not affect HIV-1 Gag expression from transfected HIV-1 proviral DNA where Tat is present (21). Conversely, Tax transactivation occurs through the mediation of interactions with host factors, specifically the CREB/CBP/p300 complex (44, 59, 60). Whether SAMHD1 interacts with host proteins to further suppress HTLV-1 LTR activity through disruption of Tax activity is unknown. As a simple retrovirus, MLV does not encode transactivation accessory proteins; however, its LTR has several binding sites for transcription factors, including nuclear factor 1 (61). Our data suggests that SAMHD1-mediated suppression of LTR activity may be specific for complex human retroviruses and could be influenced by certain transcription factors that bind to each respective LTR.

SAMHD1 enzyme activity can be regulated by mutations to its catalytic core or by post-translational modifications (62). Thus, we used two SAMHD1 mutants to determine the contribution of phosphorylation and dNTPase activity to the suppression of LTR-driven gene expression. The nonphosphorylated T592A mutant had reduced ability to suppress HIV-1 LTR-driven gene expression. Additionally, the dNTPase-inactive HD/RN mutant did not efficiently suppress HIV-1 LTR-driven gene expression. It is unlikely that a reduction in dNTP levels is required for the effect on LTR-driven gene expression as dNTP levels are high in HEK293T cells despite SAMHD1 overexpression (63). Interestingly, whereas WT SAMHD1 was observed to bind specifically to the HIV-1 LTR both in ChIP-qPCR and *in vitro* binding experiments, the SAMHD1 mutants failed to bind to the HIV-1 DNA regions tested. It is possible that SAMHD1 oligomerization may play a role in the ability of SAMHD1 to bind DNA. Dimeric SAMHD1 binds ssNA; however, previous reports have shown that tetramerization of SAMHD1 inhibits NA binding (18, 20). Phosphorylation of SAMHD1 at residue T592 destabilizes tetramer formation and impairs the dNTPase activity of SAMHD1 (64, 65), with binding of phosphomimetic T592E to ssRNA and ssDNA being identical to WT SAMHD1 (20). *In vitro,* the HD/RN mutant tetramerizes to a greater extent than WT SAMHD1 (66), and mutations of either H206 or D207 residues result in loss of ssDNA binding (15). It is possible that the T592A and HD/RN mutants form more stable tetramers and, as a consequence, lose the ability to bind the LTR and suppress activation. However, experiments to determine the oligomeric states of WT, T592A, and HD/RN SAMHD1 in the presence of fragments of the HIV-1 LTR can help to further test this possibility. Future studies are required to map the region of interaction between SAMHD1 and the HIV-1 LTR and to examine the contribution of binding to the suppression of LTR-driven gene expression. Together, our data suggests a mechanism for suppression of LTR activity in which WT SAMHD1 is able to bind directly to the LTR and possibly occlude transcription factors required for LTR activity.

As suppression of the HIV-1 LTR is a common mechanism contributing to latency (52), we aimed to determine whether SAMHD1 affects latency reversal in cells. We utilized two HIV-1 latency cell models, the J-Lat cell line and primary T_CM_ cells (40, 45, 54, 55, 67). In both models, SAMHD1 expression resulted in a suppression of latency reactivation. The modest effect observed could be due to saturation of the NF-κB pathway by PMA and TNFα (Fig. 4) or anti-CD3/CD28 (Fig. 7), as significant activation of the LTR may mask the suppressive effect of SAMHD1 (46, 50, 68, 69).

In summary, our data indicate a correlation between SAMHD1 binding to the HIV-1 LTR and SAMHD1-mediated suppression of viral gene expression and reactivation of HIV-1 latency, suggesting that SAMHD1 is among the host proteins involved in the transcriptional regulation of proviral DNA. Our results further implicate that the T592 and H206/D207 residues of SAMHD1 are important for LTR binding and suppression of HIV-1 gene expression. Future studies using latently infected cells from HIV-1 patients and primary HIV-1 isolates will further inform the function and mechanisms of SAMHD1 as a novel modulator of HIV-1 latency.

## MATERIALS AND METHODS

### Cell culture

Human embryonic kidney 293T (HEK293T) cells were obtained from the American Type Culture Collection (ATCC) and maintained as described (27). Jurkat cell-derived J-Lat cells (clone 9.2) were obtained from the NIH AIDS reagent program and maintained as described (45). THP-1 control cells and derived SAMHD1 KO cells were maintained as described (29). All cell lines tested negative for mycoplasma contamination using a PCR-based universal mycoplasma detection kit (ATCC, #30-101-2k). Healthy human donors’ peripheral blood mononuclear cells (PBMCs) were isolated from the buffy coat as previously described (70). Naïve CD4^+^ T-cells were isolated from PBMCs by MACS microbread-negative sorting and the naïve CD4^+^ T-cell isolation kit (Miltenyi Biotec). Primary CD4^+^ T-cells were cultured in complete RPMI-1640 media in the presence of 30 IU/mL of recombinant interleukin 2 (rIL-2) (Obtained from the NIH AIDS Research and Reference Reagent Program, catalog number 136) (27).

### Plasmids

The pLenti-puro vectors encoding hemagglutinin (HA)-tagged WT SAMHD1 (driven by the CMV immediate-early promoter) and the empty vector were described (4) and provided by Nathaniel Landau (New York University). The pLenti-puro vector expressing HA-tagged T592A and HD/RN SAMHD1 mutant constructs were generated using a Quikchange mutagenesis kit (Agilent Technologies) (27). The HTLV-1-LTR luciferase reporter plasmid and pcTax were provided by Patrick Green (The Ohio State University) (71). The HIV-1 FF-luc (pGL3-LTR-luc) was provided by Jian-Hua Wang (Pasteur Institute of Shanghai) (55). The pRenilla-TK plasmid was provided by Kathleen Boris-Lawrie (University of Minnesota). The pTat plasmid is a pcDNA3-based HIV-1 Tat expression construct (72) provided by Vineet KewalRamani (National Cancer Institute). The MLV-LTR reporter (pFB-Luc) contains the MLV 5’ LTR, truncated *gag*, 3’ LTR, and *firefly luciferase*, which was provided by Vineet KewalRamani (National Cancer Institute).

### Transfection assays

HEK293T cells (5.0 × 10^4^ in experiments of Fig. 1 and 1.0 × 10^5^ in experiments of Fig. 2-3) were co-transfected with a viral LTR-driven luciferase construct (HIV-1, HTLV-1, or MLV), TK-driven Renilla luciferase construct, and increasing amounts of SAMHD1 WT, T592A or HD/RN mutant-expressing plasmid using calcium phosphate as described (27). The total amount of DNA transfected was maintained through addition of empty vector. Transfection media was replaced with fresh media at 16 h after transfection. Nucleofection of control and SAMHD1 KO THP-1 cells with HIV-1-LTR-Luc and TK-Renilla was performed using the Amaxa Cell Line Nucleofector Kit V (Lonza).

### Immunoblotting and antibodies

Cells were harvested 24 h after transfection or as specifically indicated, washed with phosphate-buffered saline (PBS) and lysed with cell lysis buffer (Cell Signaling, #9803) containing protease inhibitor cocktail (Sigma-Aldrich P8340). Cell lysates were prepared for immunoblotting as described (27). HA-tagged SAMHD1 and endogenous glyceraldehyde-3-phosphate dehydrogenase (GAPDH) were detected using antibodies specific to HA (Covance, Ha.11 clone 16B12) at a 1:1,000 dilution, and GAPDH (BioRad, AHP1628) at a 1:3,000 dilution, respectively. Polyclonal SAMHD1-specific antibodies (Abcam, ab67820) were used at a 1:1,000 dilution for immunoblotting, as described (63). Immunoblots were imaged and analyzed using the Amersham imager 600 (GE Healthcare). Validation for all antibodies is provided on the manufacturers’ websites.

### Densitometry quantification of immunoblots

Densitometry analysis was performed on unaltered low-exposure images using the ImageJ software. Densitometry values were normalized to GAPDH.

### Protein expression and purification

Full-length cDNA of WT, T592A, and HD/RN SAMHD1 were cloned into a pET28a expression vector with a 6 × His-tag at the N- terminus and expressed in *E. coli*. SAMHD1 proteins were purified using a nickel-nitrilotriacetic acid affinity column as described (73). The eluted peak fractions were collected and dialyzed into the assay buffer, and then further purified with size-exclusion chromatography as described (21). SAMHD1 protein was stored in buffer containing 50 mM Tris-HCL (pH 8.0), 150 mM NaCl, 5 mM MgCl_2_, 0.5 mM Tris-(2-carboxyethyl) phosphine at −80 °C.

### Synthetic DNA oligonucleotides

Oligonucleotides used in FA binding assays, and as primers for qPCR, were synthesized by Integrated DNA Technologies. Sequences of oligonucleotides and primers are shown in Tables 1 and 2. A 90-mer 6-FAM-labeled DNA oligonucleotide derived from the scrambled sequence of the HIV-1 LTR was obtained using the Sequence Manipulation Suite (Bioinformatics.org).

### FA binding assays

The assays were performed as described (53, 74) using 5’-6-FAM-labeled DNA sequences shown in Table 2. Briefly, proteins were incubated with 10 nM DNA at room temperature for 30 min in 20 mM Tris-HCl, pH 8, 1 mM MgCl_2_, 0.25 mM HEPES, 50 μM 2-mercaptoethanol, and 50, 100, or 150 mM monovalent ions (25, 50, or 75 mM of each NaCl and KCl). Each measurement was performed in triplicate over a range of WT or mutant SAMHD1 (5-8,300 nM). Binding affinities were calculated by fitting the data to a 1:1 binding model, as described (75). Fluorescence measurements were obtained using a SpectraMax M5 plate reader (Molecular Devices, Sunnyvale, CA).

### Generation of SAMHD1-expressing J-Lat cell lines

HEK293T cells were transfected with pLenti-puro vector or HA-tagged SAMHD1 (WT, T592A, and HD/RN) expressing plasmids, pMDL packaging construct, pVSV-G, and pRSV-rev to produce lentiviral stocks for spinoculation at 2,000 × *g* for 2 h at room temperature. Lentiviral stocks were harvested, filtered, and concentrated through a sucrose cushion at 48 h post transfection. Concentrated lentivirus stock was resuspended in RPMI-1640 media and applied to J-Lat cells (clone 9.2) with polybrene (8 μg/mL) prior to spinoculation. Afterwards, cells were cultured in complete RPMI media for 72 h before undergoing selection with 0.8 μg/mL puromycin.

### HIV-1 reactivation assay in J-Lat cells

J-Lat cells (clone 9.2) stably expressing WT, mutant T592A or HD/RN SAMHD1 were generated as described above. Cells were treated with 10 ng/mL TNFα, or 32 nM PMA with 1 μM ionomycin (2× PMA+i) unless otherwise described in figure legends. At 24 h post-treatment, media was removed and cells were washed and placed in untreated complete RPMI-1640 media for an additional 12 h. Cells were collected, washed twice with 1× PBS, and suspended in 2% fetal bovine serum in PBS. Cells were evaluated by flow cytometry using Guava EasyCyte Mini Flow Cytometer (Millipore), with data analyzed by FlowJo software.

### IP of SAMHD1 in J-Lat cells

J-Lat cells (clone 9.2) expressing WT or mutant SAMHD1 and the vector control cells were differentiated using either 64 nM PMA (WT) or 128 nM PMA (vector, T592A, and HD/RN) for 24 h. At 36 h post-treatment, cells were treated with 1% paraformaldehyde for 10 min before the reaction was quenched with 0.125 M glycine. Cells were lysed in non-SDS containing radioimmunoprecipitation assay buffer and sonicated to shear cellular chromatin. Monoclonal anti-HA-agarose beads were incubated with 250 μg of cell lysate from SAMHD1-expressing (WT, T592A, HD/RN) or vector control cells at 4°C for 2 h. Beads were washed 3 times with PBS containing 0.1% Tween. To confirm IP efficiency, bound proteins were eluted from beads by boiling in 1× SDS-sample buffer, and the supernatants were analyzed by immunoblot as described (27). Total DNA was isolated from proteinase-K treated sonicated input lysate and IP products using a DNeasy kit (Qiagen).

### qPCR assay

For quantification of *Renilla* or *firefly luciferase* mRNA in transfected HEK293T cells, total cellular RNA was extracted using the RNeasy mini kit (Qiagen). Equal amounts of total RNA from each sample were used as a template for first-strand cDNA synthesis using Superscript III first-strand synthesis system and oligo (dT) primers (Thermo Fisher Scientific). SYBR green-based PCR analysis was performed using specific primers detailed in Table 1 and methods described (63). Quantification of spliced *GAPDH* mRNA was used for normalization as described (63). Calculation of relative gene expression was performed using the 2^-ΔΔCT^ method as described (76).

The levels of SAMHD1-bound HIV-1 genomic DNA from PMA-treated latently infected J-Lat cells were measured by SYBR-green-based qPCR using primers detailed in Table 1 and methods as described (63, 77). DNA samples without primer templates were used as negative controls. Genomic DNA (50 ng) from PMA-treated SAMHD1-expressing or vector control J-Lat cells after IP was used as input for the detection of HIV-1 genes. Data was normalized to vector background levels and presented as percent of total input DNA.

### Generation of shRNA vectors

HEK293T cells were transfected with pLKO.1-puro empty vector (GE DHarmacon) and SAMHD1-specific shRNA expressing plasmids (GE DHarmacon, clone ID: TRCN0000145408), psPAX2 packaging construct, and vesicular stomatitis virus G-protein-expressing construct (pVSV-G) to produce lentiviral stocks for spinoculation at 2,000 × *g* for 2 h at room temperature. Lentiviral stocks were harvested, filtered, and concentrated through a sucrose cushion at 48 h post transfection. Concentrated lentivirus stock was resuspended in RPMI-1640 media and applied to isolated naïve CD4^+^ T-cells with polybrene (8 μg/mL) prior to spinoculation. Afterwards, cells were cultured in complete RPMI-1640 media with 30 IU/mL of IL-2.

### HIV-1 latency reactivation assay in primary T_CM_ cells

We utilized the primary T_CM_ model of latency as described (54, 55). In brief, naïve CD4^+^ T-cells cells were stimulated for 72 h with anti-CD3/CD28-antibody coated magnetic beads (Dynabeads). After an additional 4 days of culture, cells were infected with VSV-G-pseudotyped HIV-1-GFP (40) and cultured for 7 days to produce latently infected T_CM_. Next, cells were transduced with lentiviral vectors containing vector control or SAMHD1 shRNA for 3 days before activation with or without anti-CD3/CD28 antibody-coated magnetic beads for 3 days. HIV-1 reactivation was measured by flow cytometry.

## ACKNOWLEDGMENTS

We thank Kathleen Boris-Lawrie, Patrick Green, Vineet KewalRamani, Baek Kim, Nathaniel Landau, Vicente Planelles, and Jian-Hua Wang for sharing reagents, and members of the Wu and Musier-Forsyth labs for valuable discussions. The following reagents were obtained through the NIH AIDS Reagent Program, Division of AIDS, NIAID, NIH: J-Lat Full Length GFP Cells (clone 9.2) from Dr. Eric Verdin, and human rIL-2 from Dr. Maurice Gately, Hoffmann - La Roche Inc.

This work was supported by NIH grants R01 AI104483, R01 AI120209, and R01 GM128212 to L.W., NIH grant R01 GM113887 to K.M.-F., C. Glenn Barber funds from the College of Veterinary Medicine at the Ohio State University (OSU) to J.M.A., NIH F31 GM119178 and a fellowship from the Center for RNA Biology at OSU to A.A.D., NIH T32 GM007223 and the National Science Foundation Graduate Research Fellowship to O.B., and NIH T32 GM008283 to K.M.K.

L.W. conceived the study and designed experiments with J.M.A. and C.S.G. J.M.A. S.H.K. and S.B. performed the experiments. O.B. and K.M.K. purified recombinant SAMHD1 proteins. J.M.A., C.S.G, A.A.D., K.M.F., Y.X., C.S.G., and L.W. analyzed data. J.M.A. and L.W. wrote the manuscript. All authors reviewed the results, revised manuscript, and approved the final version of the manuscript. All the authors declare that there are no conflicts of interest.

